# The X-ray crystal structure of BorF, the flavin reductase subunit of a two-component flavin-dependent tryptophan halogenase

**DOI:** 10.1101/2025.08.09.669344

**Authors:** Zheng Ma, Emily W. Rady, Aravinda J. de Silva, John J. Bellizzi

**Affiliations:** Department of Chemistry and Biochemistry, College of Natural Sciences and Mathematics, The University of Toledo, 2801 W. Bancroft St Toledo, OH, 43606, U.S.A

**Keywords:** Flavin adenine dinucleotide, oxidoreductase, crystallography, biosynthesis

## Abstract

BorF is a short-chain flavin reductase from a desert soil bacterium that uses NADH to reduce FAD to FADH_2_, which is used by the tryptophan-6-halogenase BorH to chlorinate tryptophan in the biosynthetic pathway of borregomycin A. The X-ray crystal structure of BorF bound to FAD was solved to 2.37 Å by molecular replacement and consists of a homodimer of single-domain protomers with a Greek key split β-barrel topology containing a domain-swapped N-terminal α-helix, as seen in other members of this family. Insertions and deletions in the region between α3 and β5 lead to a variety of different conformations of the adenosine portion of FAD bound to BorF and structurally related reductases. Comparison of the FAD-bound structures of BorF and BorH suggests that FAD must completely dissociate from BorH in order to be reduced by BorF.

## 1. Introduction

Flavin-dependent halogenases (FDHs) use FADH_2_, O_2,_ and Cl^-^ to generate HOCl, which travels down a 10 Å tunnel and halogenates an aromatic substrate in an electrophilic aromatic substitution reaction^1-4^. These bacterial and fungal biosynthetic enzymes have desirable properties for green chemistry synthetic applications due to their ability to regioselectively synthesize aryl halides under mild reaction conditions, making them valuable for the atom-efficient production of synthetically useful intermediates for cross-coupling reactions as well as biologically active halogenated target compounds^5-7^. In the process of generating HOCl, FADH_2_ is oxidized to FAD, but the FDH cannot reduce the FAD to complete the catalytic cycle; this function is “outsourced” to a flavin reductase (FR) partner protein to complete their catalytic cycle. FDH/FR pairs are a subclass of the two-component flavin-dependent monooxygenase systems, in which flavin acts as a cosubstrate/product for the two proteins, diffusing between the larger FDH subunit (which accepts FADH_2_ as a substrate and releases FAD as a product) and the smaller FR subunit (which reduces FAD to FADH_2_ using NADH)^8^.

Borregomycin A (Figure 1A) is a chlorinated bisindole alkaloid synthesized via a biosynthetic pathway encoded by a gene cluster from a soil bacterial metagenome isolated from Anza-Borrego desert soil samples^9^. Work on structurally related indolotryptoline natural products has established that their structural core is assembled via oxidative dimerization of tryptophan, and the chlorine is incorporated before dimerization by chlorination of tryptophan^10, 11^. The biosynthetic gene cluster for borregomycin A includes two genes (*borH* and *borF*) encoding proteins with homology to the halogenase and flavin reductase subunits of a two-component FDH/FR system^9^. We have shown that BorH chlorinates and brominates tryptophan at C6 *in vitro*, and can also halogenate other aromatic compounds containing indole, benzene, and quinoline groups^12, 13^. BorF and NADH are required for BorH to halogenate Trp, implicating BorF as the FR that partners with BorH (Figure 1A)^14^. BorH and BorF exhibit enhanced thermal stability (*T*_*m*_ of 48 °C for BorH and 50 °C for BorF), reflecting their origins from desert soil bacteria.

**Figure 1:**
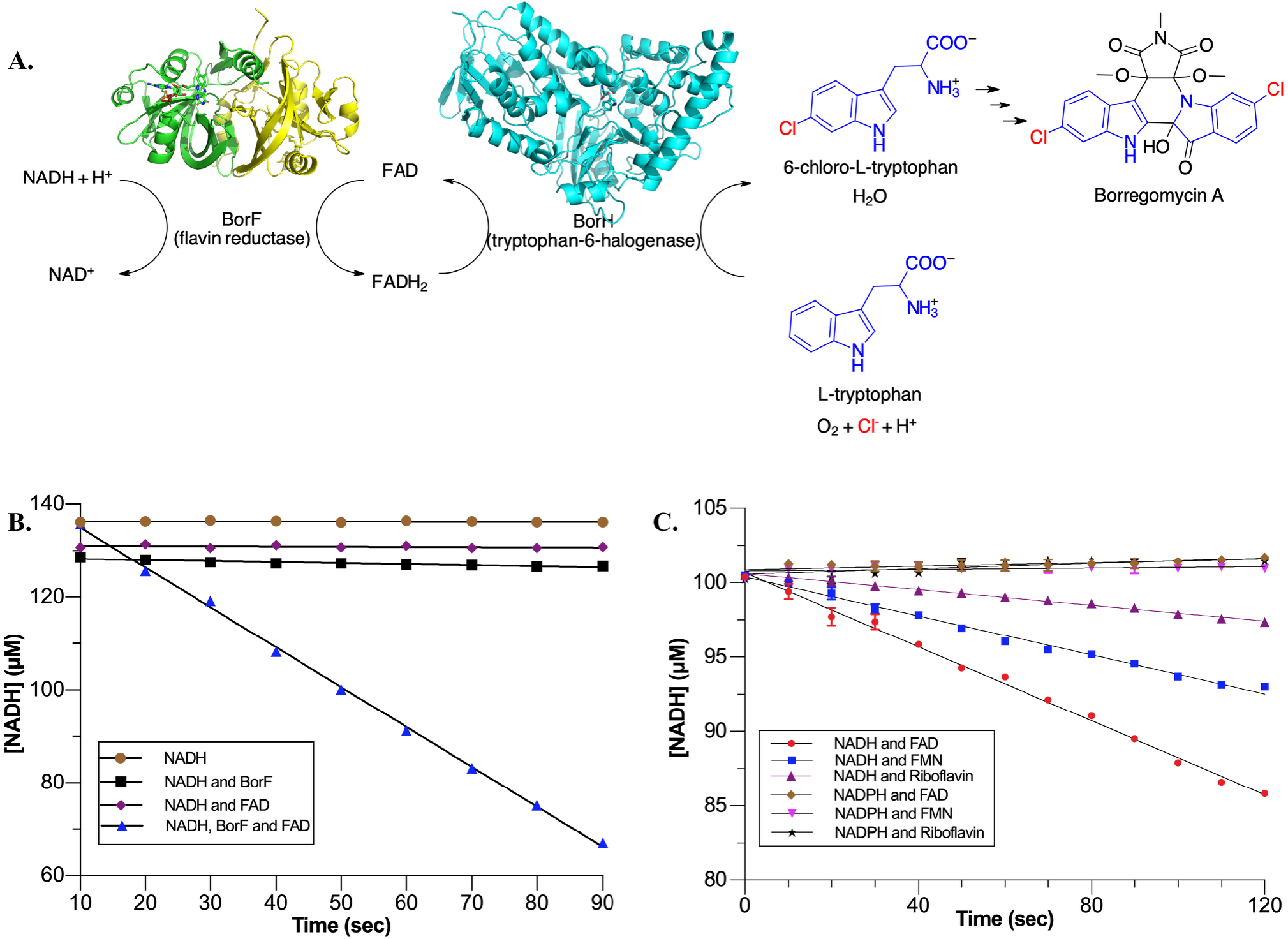
BorF is a flavin reductase. **A**. BorF supplies reduced flavin to BorH, which chlorinates Trp in the biosynthetic pathway producing borregomycin A. **B**. Oxidation of NADH (monitored by absorbance at 340 nm) only occurs in the presence of both FAD and BorF. **C**. BorF requires NADH (not NADPH) as reductant and prefers FAD over FMN and riboflavin.

BorF shares sequence homology with FRs that supply reduced flavin to two-component monooxygenases and belongs to a family of short-chain oxidoreductases (InterPro IPR002563; Pfam PF01613; SMART SM00903), which also includes enzymes that reduce cobalamin and iron. Previous crystal structures of reductases in this family^15-23^ have illuminated the flavin binding site (with bound FAD in some cases and FMN in others), and some have had nicotinamide cosubstrate/coproduct bound with the flavin in a ternary complex. These enzymes lack a Rossmann fold, binding the nicotinamide instead in a folded conformation with the nicotinamide ring sandwiched between the isoalloxazine ring of the flavin (facilitating hydride transfer) and the adenine ring. To date, the only published crystal structure of an FR known to provide FADH_2_ to a halogenase is SgcE6, part of the biosynthetic pathway producing an enediyne antitumor antibiotic, which has been shown to supply free reduced flavin to both a monooxygenase and halogenase in that pathway, both of which modify an acyl carrier protein-linked tyrosine substrate^22, 24^.

Previous work on flavin reductases from two-component flavin-dependent monooxygenase and halogenase systems has revealed diversity in kinetic mechanism and mode of flavin transfer despite high sequence and structural homology. For example, BorF uses an ordered sequential kinetic mechanism with FAD binding before NADH, whereas AbeF uses a random sequential mechanism, and PheA2 uses a ping-pong bisubstrate mechanism involving a cofactor FAD as well as a substrate FAD^14, 15^. It has been proposed that some two-component monooxygenase systems have evolved mechanisms to efficiently shuttle the flavin back and forth between the monooxygenase and reductase subunits, including protein-protein interactions, allosteric regulation of reduced flavin dissociation from the reductase, and the formation of a transient FAD-bridged flavin transfer complex^21, 25-28^. These systems may have evolved to improve efficiency, since freely diffusing FADH_2_ may react with O_2_ or other oxidants in solution. As a result, even though *in vitro* halogenation of substrates can be accomplished by FDHs using any source of FADH_2_, it has been postulated that the efficiency of the flavin transfer may be enhanced by using the specific FR that co-evolved with the FDH, though potential mechanisms of flavin transfer between the subunits have yet to be rigorously investigated. To further our understanding of the BorH-BorF two-component FDH/FR system, we have determined the crystal structure of BorF complexed with FAD.

## 2. Materials and methods

### 2.1 Macromolecule production

An expression construct encoding His_6_-MBP-BorF was produced by PCR amplification of a synthetic *borF* gene (ThermoFisher GeneArt), followed by Gibson assembly cloning (New England Biolabs) into vector p28-His_G,_ which encodes an N-terminal His_6_-maltose binding protein fusion tag with a PreScission protease cleavage site. His_6_-MBP-BorF was overexpressed in *Escherichia coli* Rosetta2 (DE3) pLysS cells (Novagen) in Luria Broth (Research Products International) with 50 μg/mL kanamycin and 30 μg/mL chloramphenicol at 37 °C with shaking until an OD_600_ of 0.6 was reached, at which point the culture temperature was lowered to 16 °C and expression was induced with 100 μM IPTG. After 20 hours post-induction growth at 16 °C, cells were harvested by centrifugation at 12,000 x g, resuspended in 20 mM Tris pH 7.5, 500 mM NaCl, 1 mM TCEP, with an EDTA-free protease inhibitor tablet (Roche), and lysed by sonication. His_6_-MBP-BorF was captured from the clarified lysate by affinity chromatography using an MBPTrap HP 5 mL column on an AktaPurifier 10 system (GE Healthcare). Pooled His_6_-MBP-BorF fractions were treated with PreScission protease at 4 °C for 16 hours to cleave the His_6_-MBP from BorF, and the protease and cleaved His_6_-MBP were removed using Co (II)-charged TALON IMAC resin (Clontech). BorF was further purified using a Superdex 200 10/300 GL size exclusion column (GE Healthcare) in 20 mM Tris pH 7.5, 150 mM NaCl. BorF fractions are yellow in color, indicating copurification with flavin. This flavin can be removed while the protein is immobilized on the affinity column using 2 M KBr and 2 M urea^29^, and mass spectrometry of the removed flavin confirmed that the flavin removed was FAD, not FMN. BorF copurified with flavin (holo-BorF) was used for crystallization, and BorF stripped of flavin (apo-BorF) was used for activity studies.

### 2.2 Flavin reductase activity

Flavin reductase activity was determined by measuring depletion of NADH spectrophotometrically at 340 nm. Room temperature reactions were prepared containing 25 mM HEPES pH 7.5, 125-140 μM NADH, and either no enzyme, 0.2 μM apo-BorF, or 0.2 μM apo-BorF + 25 μM FAD in a total reaction volume of 200 μL. Absorbance measurements were recorded using Molecular Devices SpectraMax microplate reader and converted to concentrations using the extinction coefficient for NADH (6220 M^-1^ cm^-1^). To study substrate specificity, reactions were carried out with all six permutations of three hydride acceptors (25 μM FMN, riboflavin, or FAD), and two hydride donors (100 μM NADPH or NADH).

### 2.3 Crystallization

Purified holo-BorF (10 mg/mL, 0.474 mM, in 20 mM Tris pH 7.5 100 mM NaCl) was incubated with 1 mM FAD (MP Biomedical) for co-crystallization of the BorF/FAD complex. Crystals were grown using the hanging drop vapor diffusion method, in which 1 μL of BorF/FAD solution was mixed with an equal volume of reservoir solution containing 0.1 M HEPES pH 7.6, 20% (*v/v*) polyethylene glycol (PEG) 400 and 5% (*v/v*) glycerol and equilibrated at room temperature against 200 μL of reservoir. Yellow crystals appeared after 2 days at room temperature and grew to overall dimensions of 0.03 × 0.03 × 0.025 mm. Crystals were cryoprotected in mother liquor containing 25% (*v/v*) PEG 400 and 5% (*v/v*) glycerol, mounted and flash-cooled in liquid N_2_.

### 2.4 Data collection and processing

Diffraction data were collected at a wavelength of 0.98757 Å and a temperature of 100 K using the oscillation method at the Life Sciences Collaborative Access Team beamline 21-ID-G at the Advanced Photon Source (Argonne National Laboratory). Reflections were indexed and integrated using *imosflm*^30-32^, scaled and merged with *Scala*^33^, and converted to structure factor amplitudes using *Truncate*^34^. The high-resolution cutoff used for model building and refinement was 2.37 Å, reflecting the highest resolution shell with an average I/σ(I) > 2.0 (Table 1).

**Table 1:** Data collection and refinement data. Values for the outer shell are given in parentheses.

|  |  |
| --- | --- |
| Diffraction source | APS beamline 21-ID-G |
| Wavelength (Å) | 0.98757 Å |
| Temperature (K) | 100 |
| Detector | Marmosaic 300 mm CCD |
| Space group | *C*2 |
| *a*, *b*, *c* (Å) | 278.01, 62.34, 101.88 |
| $\alpha$, $\beta$, $\gamma$ (°) | 90, 110.84, 90 |
| Resolution range (Å) | 47.61–2.37 (2.45–2.37) |
| Total No. of reflections | 306720 (30373) |
| No. of unique reflections | 66699 (6590) |
| Completeness (%) | 99.93 (99.97) |
| Redundancy | 4.6 (4.6) |
| $\langle I/\sigma(I) \rangle$ | 9.4 (2.0) |
| Overall *B* factor from Wilson plot (Å<sup>2</sup>) | 44.21 |
| *R*<sub>merge</sub> | 0.100 (0.820) |
| *R*<sub>meas</sub> | 0.113 (0.923) |
| CC<sub>1/2</sub> | 0.995 (0.750) |
| Reflections used in refinement | 66699(6590) |
| Reflections used for *R*<sub>free</sub> | 1378 (129) |
| Final *R*<sub>work</sub> | 0.215 (0.296) |
| Final *R*<sub>free</sub> | 0.247 (0.351) |
| No. of non-H atoms |  |
| Protein | 9688 |
| Ligand (FAD) | 424 |
| Water | 157 |
| Total | 10269 |
| R.m.s. deviations |  |
| Bonds (Å) | 0.004 |
| Angles (°) | 0.990 |
| Clashscore | 2.20 |
| Average *B* factors (Å<sup>2</sup>) |  |
| Protein | 53.5 |
| Ligand | 52.6 |
| Water | 39.1 |
| Ramachandran plot |  |
| Most favored (%) | 98.67 |
| Allowed (%) | 1.33 |

### 2.5 Structure determination and refinement

Molecular replacement was performed with *Phaser-MR*^35^ using chain A of PDB entry 4HX6^22^ (SgcE6; 42% sequence identity to BorF) as a search model. *Phenix*^36^ was used for model building and refinement. Eight chains arranged as four homodimers were built within the asymmetric unit using the *Phenix* autobuild module^37^. After rigid body refinement using *Phenix*.*refine*^38^, electron density for FAD was clearly visible for all eight subunits in αA-weighted *F*_*o*_*-F*_*c*_ maps. The *LigandFit*^39-41^ and *elBOW*^39^ modules of *PHENIX* were used to fit FAD to the density. Iterative model building using *Coot*^42^ and positional, real space, simulated annealing, and individual isotropic B-factor refinement using *Phenix*.*refine* converged to a final model with an *R*_*work*_ of 21.9% and an *R*_*free*_ of 24.6%. Torsion-based NCS restraints were used in the early rounds of refinement, and water picking and deletion was carried out automatically with *Phenix* followed by manual inspection of water density and hydrogen bonding using *Coot*. X-ray/stereochemistry and X-ray/ADP weights were optimized during the final round of refinement. Data collection and refinement statistics are presented in Table 1. *MolProbity*^43^ was used to validate the final model. DALI^44^ was used to identify structural homologs. Figures were produced using Endscript/ESPript^45^, PyMOL^46^, and Ligplot^+47^. Model coordinates and structure factors were deposited in the RCSB Protein Data Bank with PDB code 5CHO.

## 3. Results and discussion

### 3.1 BorF is a flavin reductase that requires NADH and prefers FAD

BorF oxidizes NADH to NAD^+^, as determined by NADH depletion monitored by absorbance at 340 nm. NADH oxidation requires the presence of both FAD and BorF (Figure 1B). BorF prefers FAD as a substrate but is also able to reduce FMN and riboflavin; however, it is incapable of using NADPH as a hydride source. (Figure 1C).

### 3.2 Structure of the BorF/FAD complex

The structure of the BorF/FAD complex was solved at 2.37 Å by molecular replacement using SgcE6 (4HX6-A; 42% identity) as a search model. BorF co-crystallized with FAD in space group *C2* with four homodimers (AB, CD, EF, GH) in the asymmetric unit. The crystallized construct comprises full-length BorF (amino acid residues 1-196) with an additional N-terminal proline remaining after affinity tag cleavage. However, no electron density was observed for the N-terminal 30 amino acid residues, and density for one to four residues at the C-terminus was absent in all chains. The backbone electron density for Chains A-E was clear and unbroken and side chains were well resolved except for several solvent-exposed arginine side chains, which were truncated to Cβ in the final model. Chains F-H were built into regions of the map that were noisier and contained regions of weaker electron density in loop regions, resulting in higher average temperature factors (average B factor for chains A-E = 44.1 Å^2^, average B factor for chains F-H = 73.4 Å^2^). The Cα R.M.S. deviations ranged between 0.14-0.21 Å among the eight chains in the model (Table S1). Our structural analysis will focus on the CD dimer, which has the greatest number of residues modeled and the lowest B-factors.

BorF (UNIPROT M9QXS1; GenBank AGI62216.1) belongs to a family of short-chain oxidoreductases (InterPro IPR002563; Pfam PF01613; SMART SM00903) that includes other flavin reductase components of two-component flavin-dependent monooxygenases and halogenases, as well as a wider variety of confirmed and putative oxidoreductases, including enzymes that reduce cobalamin and iron. Members of this family are homodimers with protomers containing a split β-barrel with Greek key topology, which has been described as a circular permutation of the ferredoxin reductase-like flavin binding domain^48^. Each BorF subunit comprises 11 β-strands, 3 α-helices and two 3_10_ helices (Figures 2A, 2B). The N-terminal helix α1 is domain-swapped and sits atop the β-barrel in its dimer counterpart (Figure 2C). The split β-barrel is made up of strands β1-5 and β8-9 and is capped on one end by helix α2 and on the other end by α1 from the dimer partner. Packed against one face of the barrel is a subdomain inserted between β5 and β8 that consists of η1, α3, η2, a hairpin formed by β6-β7, and the intervening loops. This insertion, along with loop β4-α2, forms a groove that cradles the ribityl, pyrophosphate and ribose groups of the FAD. C-terminal to the β-barrel, a β-hairpin formed by β10-β11 contributes to both the FAD binding site and the dimer interface.

**Figure 2:**
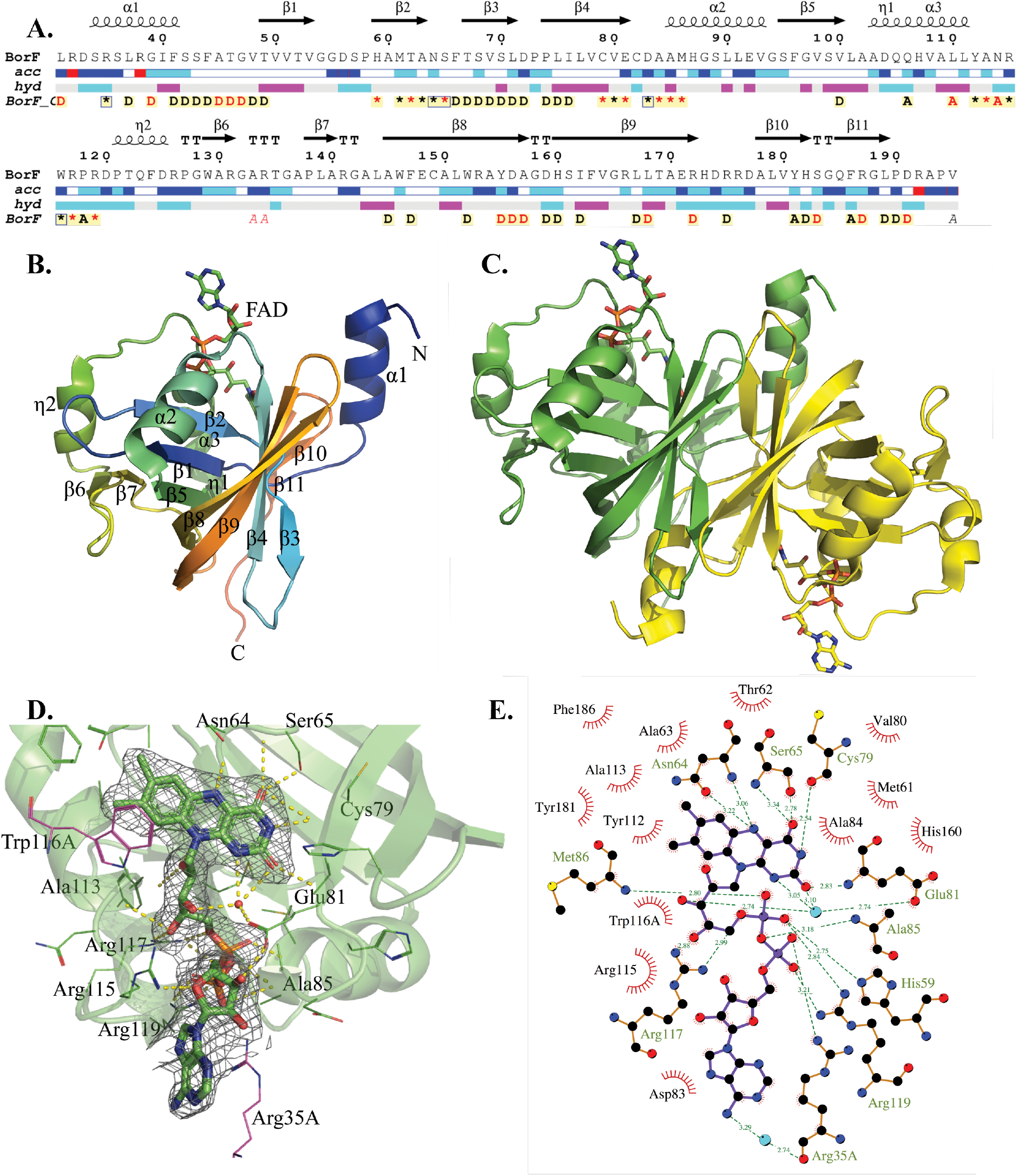
The X-ray crystal structure of BorF/FAD. **A**. BorF primary and secondary structure. Chain C of the model contains amino acids 31-195. Secondary structural elements are shown above the sequence. Relative solvent accessibility is shown below the sequence: white = buried (A < 0.1), cyan = intermediate (0.1 ≤ A ≤ 0.4), blue = accessible (0.4 ≤ A ≤1). Solvent accessibility was not calculated for Arg 32, Arg 28 and Arg 192, which were modeled without side chains (red boxes). Hydropathy is shown below the accessibility: pink=hydrophobic, grey = intermediate and cyan = hydrophilic. Intermolecular contacts are displayed below hydropathy (Red = distance < 3.2 Å and black = distance = 3.2-5.0 Å). Residues contacting Chain A (in adjacent dimer in the ASU), Chain D (dimer partner) or FAD are denoted by A, D or * respectively. **B**. Ribbon diagram of chain C of BorF/FAD with secondary structural elements labeled. FAD is shown as a stick model. **C**. BorF/FAD homodimer (chain C in yellow, chain D in green), with FAD shown as a stick model. **D**. *F*_*o*_*-F*_*c*_ Polder omit map (omitting FAD atoms) contoured at 3σ over final refined model of FAD from chain C (green stick model). Side chains contacting FAD in chain C are shown as green wire models. Side chains from chain A in adjacent dimer contacting FAD (Trp116A and Arg35A) are shown as magenta wire models. Trp116A is occupying the area occupied by NAD^+^ in the FAD/NAD^+^ ternary structures of SgcE6 (4R82), PheA2(1RZ1), and HpaC(2ED4) and is presumably the reason that soaking and cocrystallization experiments with NAD^+^ failed. **E**. Polar and nonpolar interactions between BorF and FAD. Hydrogen bonds are represented by dashed lines and hydrophobic interactions by spiked semicircles.

The dimer interface (Figure 2C) buries approximately 2300 Å^2^ and involves 60 residues from each chain. The domain swapped N-terminal helix α1 is amphipathic and inserts the side chains of Leu37C, Ile40C and Phe41C into the interior of the β-barrel, where they contact the side chains of Leu71D, Ile76D, Val100D, Ala145D, and Phe147D from the dimer partner. The linkers connecting α1 to β1 in the two protomers are antiparallel to one another in extended conformations and interact via backbone hydrogen bonds. One face of the β-barrel (β3-β4-β9-β8) interacts with its counterpart in an antiparallel fashion mediated primarily by packing of hydrophobic side chains, a π-stacking interaction (Tyr155C-Tyr155D), and hydrogen bonds between the side chain of Tyr155C and the backbone of Asp156D and between the side chain of Asp159C and the amides of Leu71D and Asp72D. Loop β2-β3 in chain C interacts with β3 in chain D through hydrophobic interactions and a hydrogen bond between Thr67C and Thr67D (at the N-terminal ends of β3C and β3D). The β10-β11 hairpin forms an antiparallel β-sandwich with its counterpart in the dimer partner with a hydrophobic core containing Val180 and Leu189 from both chains along with Cβ and Cγ of Arg187 and Cβ of His182. The side chain of Arg187C donates a hydrogen bond to Pro190D and forms a salt bridge with Asp191D, and the side chain of Arg172C forms a hydrogen bond with the side chain of Ser183D.

### 3.3 The FAD binding site

Electron density for FAD was visible in difference maps in all eight chains (Figure 2D). The binding of the isoalloxazine is consistent with previous crystal structures of short chain flavin reductases bound to FAD or FMN. The isoalloxazine occupies a pocket surrounded by strand β4, loop β2-β3, loop α1-β1, helix α3, and the β10-β11 hairpin. The 2,4-pyrimidinedione portion of the isoalloxazine forms hydrogen bonds with backbone atoms of Asn64, Ser65, Cys79 and Glu81, the side chain of Ser65, and a water molecule that also forms hydrogen bonds to the side chain of Glu81 and O3 of the ribityl group of FAD (Figure 2D, 2E). The backbone amide of Asn64 forms a hydrogen bond to FAD N5. The dimethylbenzene binds between helix α3 and the β10-β11 hairpin and contacts the side chains of Val48, Thr62, Asn64, Tyr181, Phe186, and Ala113. Trp116 from an adjacent dimer in the asymmetric unit makes a hydrophobic crystal contact with the dimethylbenzene portion of the FAD.

The ribityl group of FAD binds in an extended conformation along strand β2, hydrogen bonding with the backbones of Thr62, Tyr112, and Ala113, a water molecule, and the side chain of Arg117. The pyrophosphoryl group extends at a right angle to the ribityl group and interacts via salt bridges with the side chains of His59, Arg117, and Arg119 as well as Arg35 from an adjacent dimer in the asymmetric unit, and via hydrogen bonds with the backbones of Ala85 and Met86. The adenosine emerges from the groove at the interface between dimers in the asymmetric unit. The ribose forms hydrogen bonds with the side chains of Glu81 and Arg115. As a result of crystal packing, a cation-π interaction is formed between Arg35 from chain A (in an adjacent dimer in the asymmetric unit) and the FAD adenine group of chain C, which is approximately 4 Å away from and oriented parallel to the guanidinium group of the Arg35A side chain. The Arg35A side chain also forms a salt bridge with the one of the phosphoryl groups of FAD as noted above.

### 3.5 Trp116 from an adjacent BorF dimer obstructs the nicotinamide binding site

Some homologs of BorF have had structures determined of dead-end ternary complexes with FAD and NAD^+^ (4R82, 1RZ1, 2ED4, 2D37, 3K88). In these structures, the NAD^+^ binds in a folded conformation in which the adenine stacks on the nicotinamide ring, which in turn stacks on the central ring of the isoalloxazine group of FAD, juxtaposing C4 of the nicotinamide with N5 of the FAD for hydride transfer. We collected data on multiple BorF crystals grown in the presence of both FAD and NAD^+^ or grown in the presence of FAD and soaked with NAD^+^ but could not detect any electron density in the substrate binding site that would indicate the presence of the dead end BorF/FAD/NAD^+^ ternary complex. Crystal packing explains the lack of observed NAD^+^ binding, as the side chain of Trp116A is occupying the position where the ribose of NAD^+^ and two water molecules bridging the NAD^+^ and PheA2 are found in the PheA2 ternary complex (Figure S1).

### 3.4 Comparison of BorF to other short chain flavin reductases

DALI was used to identify structural homologs to chain C of BorF (Figure 3A and Table S2). PDB entries representing 30 different proteins with DALI Z-scores of 19.9 or higher had Cα RMSD values between 1.4-2.5 Å with BorF chain C. All are known or putative flavin-binding proteins from bacteria or archaea and can be divided into three categories. The first are enzymes known to be reductase components of two-component flavin-dependent monooxygenase systems, including SgcE6 from *Streptomyces globisporus* C-1027 (4R82/4HX6)^22^, C1-HPAH from *Acinetobacter baumannii* (5ZC2)^23^, PheA2 from *Geobacillus thermoglucosidasius* (1RZ0/1RZ1)^15^, TftC from *Burkholderia cepacian* (3K86/3K87/3K88)^19^, HpaC from *Thermus thermophilus* (2ED4/2ECU/2ECR)^17^, HpaC from *Sulfolobus tokodaii* (2D36/2D37/2D38)^16^, and SMOB from *Pseudomonas sp. Y2* (4F07)^21^. The second category are enzymes which use reduced FMN to reduce a variety of chemically distinct substrates, such as corrin reductase CobR (3CB0/4IRA) from *Brucella melitensis*^18^, ferric reductase FerA from *Paracoccus denitrificans* (4XJ2)^49^, ketoreductase PlmKR1 from *Streptomyces sp*. HK803 (4HXY)^50, 51^, ferric reductase FeR from *Archaeoglobus fulgidus* (1I0S/1I0R), YP_005476 from *Thermus thermophilus* (3ZOE, 3ZOF, 3ZOH)^52^, and 2-nitrobenzoate nitroreductase NbaA from *Pseudomonas fluorescens* (4Z85)^53^. The third set of homologs are putative flavin-binding proteins of unknown function solved by structural genomics projects, including 4L82, 2R0X, and 3RH7.

**Figure 3:**
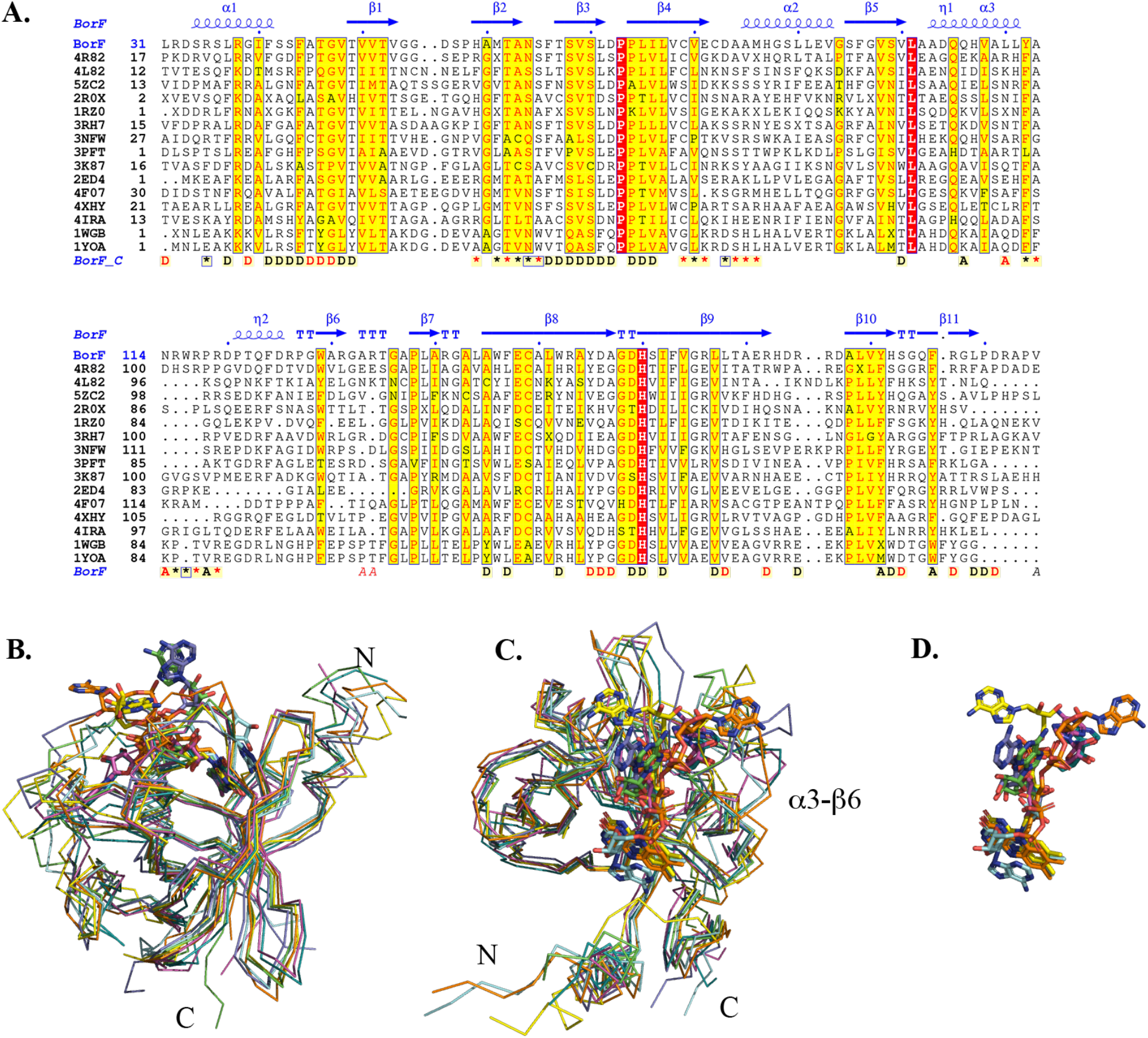
Structural homologs of BorF. **A**. Multiple sequence alignment of BorF with structural homologs (See Table S2). BorF secondary structural elements are indicated above the alignment. BorF residues contacting other subunits or FAD are labeled as in Figure 2A. Strictly conserved residues have a red background. **B**. Backbone superposition of BorF/FAD chain C (green) with selected homolog/FAD complexes, in the same orientation as Figure 2B. Green = BorF/FAD, aqua = PheA2/FAD (1RZ0), purple = SMOB/FAD(4F07), yellow = 1YOA/FAD, orange = CobR/FAD (4IRA), cyan = TftC/FAD(3K87), magenta = HpaC_TT_/FAD(2ED4). **C**. Different orientation of B, showing that the isoalloxazine and ribityl groups superimpose perfectly but there is a range of conformations of the pyrophosphate and adenine groups of FAD, which can be explained in part by differences in insertion length and secondary structure in the region between α3 and β6. **D**. Superposition of FAD from BorF and homolog complexes in same orientation as C.

The core β-barrel of BorF superimposes well with the structural homologs (Figure 3B, 3C), but the region between α3 and β5 (residues 112-129 in BorF), which forms part of the FAD binding site, varies substantially in length and sequence. The binding of the isoalloxazine portion of FAD, which primarily involves contacts with main chain atoms, is well-conserved between the BorF/FAD complex and structures of FAD and FMN bound to structural homologs of BorF. However, in structures with bound FAD, the ribityl, pyrophosphoryl, and adenylyl groups are found in a range of different conformations, primarily due to structural variation in the region between α3 and β6 (Figure 3B-3D). In BorF, α3 is followed by loop α3-η2 (residues 112 to 120), which forms hydrogen bonds and salt bridges using backbone and side chain atoms to the ribityl, pyrophosphate, and adenosine of FAD. Residues 121-125 make up 3_10_ helix η2, which is roughly antiparallel to α3, and a loop from 126-129 connects to the N-terminus of the β6-β7 hairpin. Many proteins in this structural family preferentially bind FMN rather than FAD, and the much higher conservation of the binding sites for the isoalloxazine and ribityl groups compared to the AMP moiety of FAD suggest that the fold originally evolved as an FMN-binding module but in some cases co-evolved with FAD-specific monooxygenases and halogenases, which selected for residues that make additional interactions with the AMP. In the case of SMOB, it has been suggested that the protrusion of the AMP moiety from the FR subunit may allow it to serve a bridging function between the two subunits^21, 54^.

### 3.6 The adenosine moiety of FAD is solvent-exposed and adopts variable conformations in different FRs

Of the crystal structures of BorF homologs with FAD bound, the orientation of the FAD pyrophosphoryl group and adenosine in styrene monooxygenase component 2 (SMOB/FAD; 4F07, Figure S2A) is most like BorF/FAD. Compared to BorF, SMOB has a shorter insertion between α3 and β6 and lacks the 3_10_ helix η2. A second conformation of FAD, strikingly different from that seen in BorF and SMOB, can be seen in PheA2 (1RZ0, Figure S2B) and HpaC_Tt_ (2ED4, Figure S2C). When comparing BorF/FAD to PheA2/FAD, the isoalloxazine, ribityl and first phosphoryl group superimpose well, but the second phosphoryl group in PheA2 is rotated by 180°, directing the adenosine away from the dimer interface and instead into the space between β2, α3, and loop α3-β6, where the adenine is stabilized by 2 H-bonds and a π-stacking interaction. This orientation of the adenosine is not possible in BorF, as loop α3-η2 occupies the space occupied by the adenine in PheA2/FAD. The region between α3 and β6 is shorter in PheA2 (12 amino acids compared to 17 in BorF) and lacks the 3_10_ helix η2. HpaC_Tt_ also lacks the 3_10_ helix, has a shorter β5-β6 hairpin, and has a nearly identical FAD conformation to PheA2.

Several other FRs have structural differences that sterically prevent the FAD from adopting the conformation found in the BorF structure. In the corrin reductase CobR (4IRA, Figure S2D), the isoalloxazine and C1’-C3’ of the ribose align with the FAD conformation found in BorF, but the rest of the FAD is shifted significantly due to bulkier residues in loop β4-α2 and the first turn of α2 (particularly His67 and Glu69, which are both Ala in BorF) on one side, and a different backbone conformation for loop α3-η2 that moves η2 closer to α3. Putative flavoprotein 1YOA (Figure S2E) also obstructs BorF’s adenine conformation with bulkier residues in loop β4-α2, and changes in the α3-β6 region including the deletion of η2 accommodate an alternate positioning of the pyrophosphate and ribose. FAD bound to TftC adopts two different conformations. In the ternary TftC/FAD/NAD^+^ complex (3K88), the adenine binds in an extended conformation that clashes with loop α3-η2 in BorF, but in the TftC/FAD binary complex (3K87), one protomer has the extended FAD conformation, and the other has a folded FAD with the adenine stacked with the isoalloxazine in the position occupied by the nicotinamide of NAD^+^ in the ternary complex (Figure S2F).

### 3.7 Comparison of BorF/FAD and BorH/FAD indicates that the FAD cannot simultaneously bind to BorH with its adenosine and BorF with its isoalloxazine

In BorF, the isoalloxazine of the FAD is enclosed within the active site, and the adenosine group is solvent-exposed. In contrast, in our crystal structure of BorH, the FDH partner protein of BorF, bound FAD has weaker electron density for the isoalloxazine and ribityl groups as compared to the adenosine group, suggesting the adenosine group remains tightly bound while the isoalloxazine may be mobile. This has also been observed in the tryptophan-6-halogenase Thal, leading to the speculation that rather than releasing the flavin into solution to diffuse between the two subunits, the adenosine may stay bound to the halogenase as the isoalloxazine moves back and forth between the active sites of the halogenase and the flavin reductase^55^. However, analysis of the structures of BorF/FAD and BorH/FAD (8TTI-C^13^) indicates that this would be impossible (Figure S3). C6’ of the adenosine has a maximum distance of approximately 15 Å from N6 of the isoalloxazine when FAD is in a fully extended conformation. The closest distance between C6’ of the adenosine from the BorH-bound FAD and N6 of the isoalloxazine from the BorF-bound FAD that can be achieved without steric clashes between the two proteins is approximately twice this distance (27 Å), suggesting that dissociation of the FAD adenosine from BorH is necessary for the isoalloxazine to bind the active site of BorF.

## Author Contributions

Z.M. carried out expression and purification of BorF, crystallization, X-ray diffraction data collection, molecular replacement, model building, refinement, and structural analysis. E.R performed BorF expression, purification, and crystallization. A.J.D.S. carried out expression and purification of BorF and flavin reductase activity assays. J.J.B. carried out model building and refinement, data analysis, and structural analysis. The manuscript was written by Z.M. and J.J.B.

## Acknowledgements

This research used resources of the Advanced Photon Source, a U.S. Department of Energy (DOE) Office of Science User Facility operated for the DOE Office of Science by Argonne National Laboratory under Contract No. DE-AC02-06CH11357. Use of the LS-CAT Sector 21 was supported by the Michigan Economic Development Corporation and the Michigan Technology Tri-Corridor (Grant 085P1000817). This work was supported by the National Institutes of Health (award 1R15GM144877-01 to J.J.B.). The content is solely the responsibility of the authors and does not necessarily represent the official views of the National Institutes of Health.

Authors declare no competing financial interests.

