## Supplementary Material for "The X-ray crystal structure of BorF, the flavin reductase subunit of a two-component flavin-dependent tryptophan halogenase"

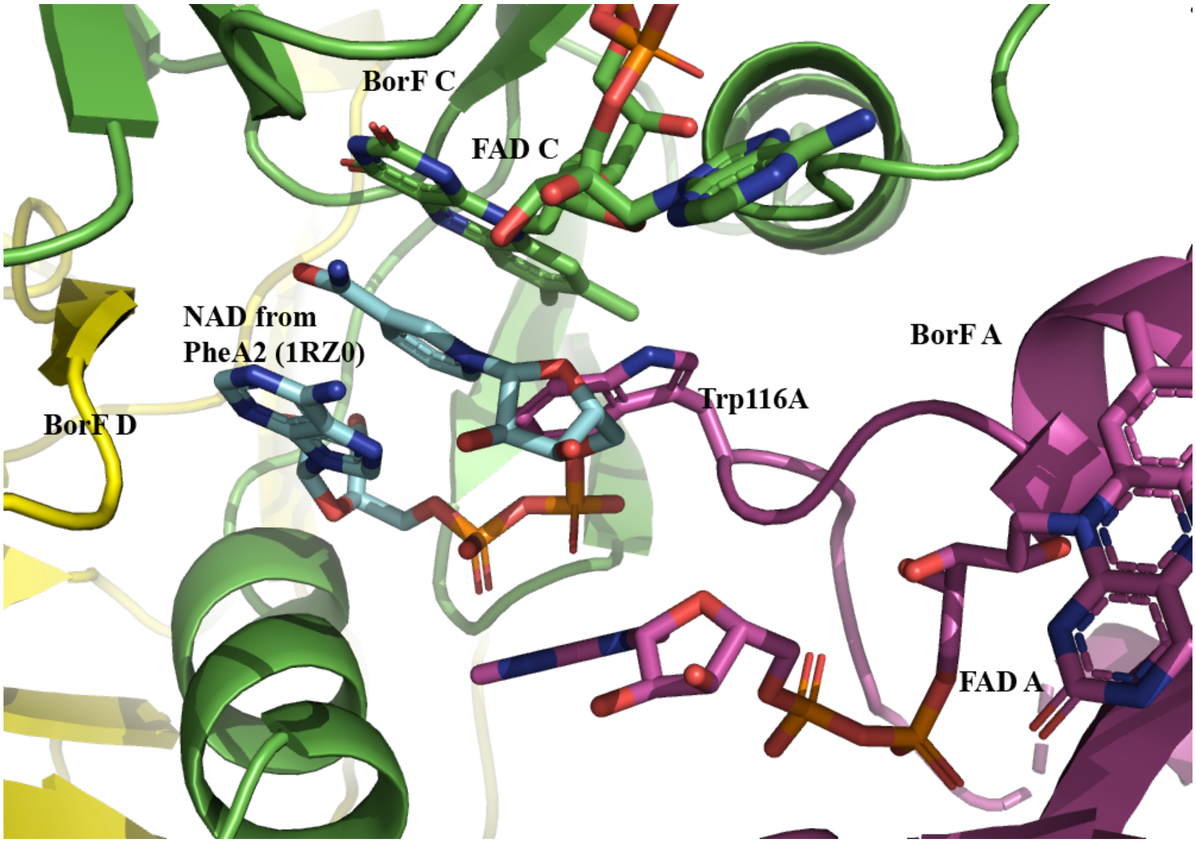


**Figure S1: BorF crystal contact obstructs $NAD^+$ binding site.**

$NAD^+$ (cyan) from chain A of the PheA2/FAD/$NAD^+$ structure (1RZ0) is shown in the active site of chain C of BorF/FAD (5CHO) after superimposing the two chains. A crystal contact from BorF chain A (in an adjacent dimer in the asymmetric unit; magenta) obstructs the $NAD^+$ binding site in BorF chain C (green). The side chain of Trp116A is occupying the position where the ribose of $NAD^+$ and two water molecules bridging the $NAD^+$ and PheA2 are found in the PheA2 ternary complex The side chain of Trp116 adenine of chain A blocks the position where the nicotinamide-connected ribose binds in the PheA2 ternary complex structure. This crystal contact would block $NAD^+$ binding and explains why BorF/FAD crystals grown in the presence of $NAD^+$ or soaked in $NAD^+$ had no electron density for $NAD^+$.

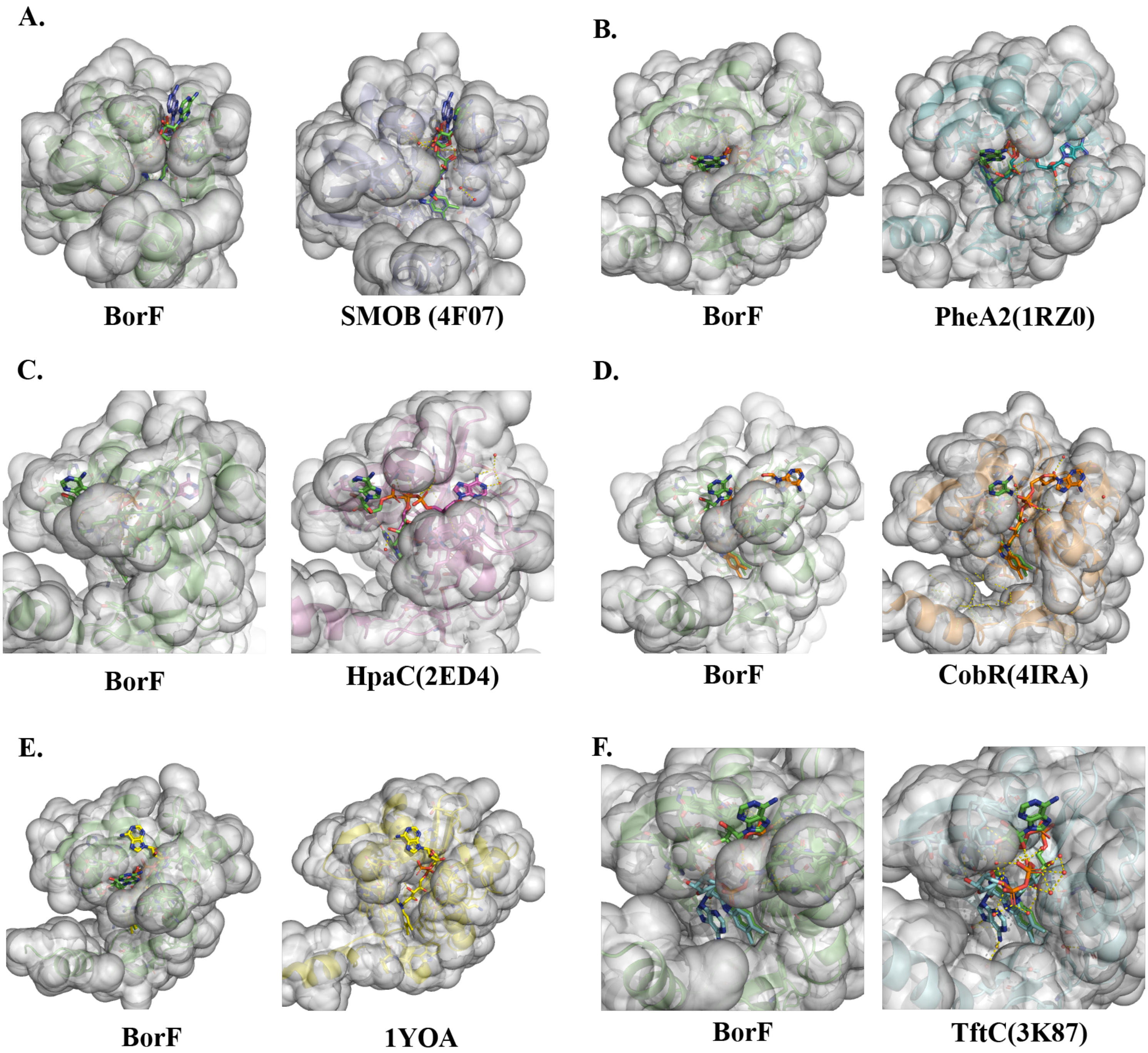


**Figure S2: Diversity of FAD binding conformations in BorF and structural homologs**.

In panels A-F, the left image shows the surface of the BorF/FAD complex with FAD from the homolog complex superimposed, and the right images shows the surface of the homolog/FAD complex with the FAD from BorF superimposed. Both images in each panel are in the same orientation as one another, but the orientation differs from panel to panel to show differences more clearly. **A.** BorF (green) and SMOB (4F07, purple). **B.** BorF and PheA2 (1RZ0, cyan) **C.** BorF and $HpaC_{Tt}$ (2ED4, magenta). **D.** BorF and CobR (4IRA, orange) **E.** BorF and 1YOA (yellow) **F.** BorF and TftC (3K87, cyan).

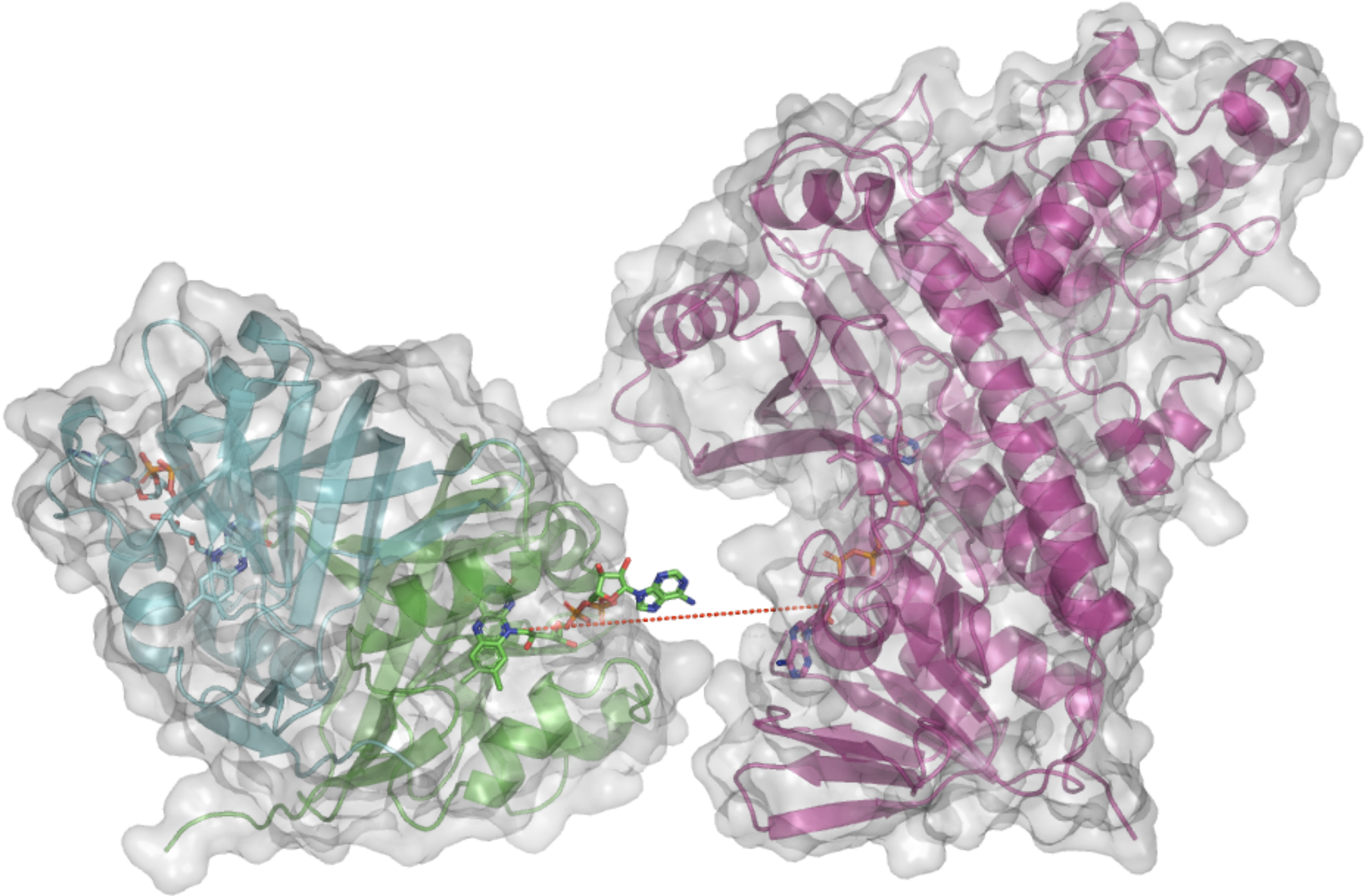

**Figure S3: BorF cannot reduce FAD that is still bound to BorH.**

Surface diagrams of BorF/FAD (5CHO chain C cyan, Chain D green) and BorH/FAD (8TTI chain C, magenta). The red dashed line connects C6’ of the adenosine from the BorH-bound FAD and N6 of the isoalloxazine from the BorF-bound FAD. The maximum distance between those two atoms in a fully extended conformation of FAD is 15 Å, and the distance shown in the figure is 27 Å. Though the isoalloxazine is buried and the adenosine is partially or completely solvent-exposed in both complexes, there is no orientation of the two proteins that would allow a single FAD molecule to simultaneously bind BorH with its adenosine and BorF with its isoalloxazine without a significant conformational change in one or both proteins. This indicates that BorF is not capable of reducing the isoalloxazine of an FAD molecule still bound to BorH via its adenosine.

**Table S1.** Pairwise backbone comparisons of BorF chains to Chain C

| Chain | Residues | Residues aligned | Cα rmsd |
|---|---|---|---|
| A | 30-194 | 165 | 0.201 |
| B | 31-193 | 163 | 0.201 |
| D | 31-194 | 164 | 0.169 |
| E | 33-193 | 161 | 0.212 |
| F | 33-192 | 160 | 0.142 |
| G | 33-192 | 160 | 0.185 |
| H | 34-193 | 160 | 0.161 |

**Table S2**. Structural homologs of BorF from the PDB

| PDB | Complex | UniProt | Protein | Gene name | Organism | DALI Z-score | Cα rmsd | Residues aligned | % sequence Identity |
|---|---|---|---|---|---|---|---|---|---|
| 4r82 | FAD/NAD | Q8GME2 | SgcE6 Flavin reductase | SGCE6 | *Streptomyces globisporus* | 25.1 | 1.4 | 156 | 48 |
| 4l82 | FMN | Q4UKE8 | Uncharacterized protein | RF_1132 | *Rickettsia felis* | 25.2 | 1.5 | 155 | 32 |
| 4hx6 | apo | Q8GME2 | SgcE6 Flavin reductase | SGCE6 | *Streptomyces globisporus* | 25.1 | 1.5 | 158 | 48 |
| 5zc2 | FMN | Q6Q271 | p-hydroxyphenylacetate 3-hydroxylase, reductase component | C1-hpah | *Acinetobacter baumannii* | 23.1 | 1.6 | 154 | 32 |
| 5zyr | FMN | Q6Q271 | p-hydroxyphenylacetate 3-hydroxylase, reductase component | C1-hpah | *Acinetobacter baumannii* | 20.3 | 1.6 | 154 | 33 |
| 2r0x | apo | Q0I3S1 | Possible flavin: NADH reductase | ycdH HS_1225 | *Haemophilus somnus* | 24 | 1.7 | 155 | 23 |
| 1rz0 | FAD | Q9LAG2 | PheA2 phenol 2-hydroxylase component B | pheA2 | *Geobacillus thermoglucosidasius* | 23 | 1.7 | 149 | 31 |
| 1rz1 | FAD/NAD | Q9LAG2 | PheA2 phenol 2-hydroxylase component B | pheA2 | *Geobacillus thermoglucosidasius* | 22.9 | 1.7 | 149 | 31 |
| 3rh7 | FMN | Q92ZM6 | Uncharacterized protein | SMa0793 | *Rhizobium meliloti* | 22.1 | 1.7 | 154 | 40 |
| 3nfw | apo | E5Q9D7 | nitrilotriacetate monooxygenase component B | NTA-Mo | *Mycolicibacterium thermoresistibile* | 21.7 | 1.7 | 156 | 29 |
| 3pft | FMN | B6CDL6 | DszD Flavin reductase (Oxidoreductase) | AFA91_09010 | *Mycobacterium goodii* | 21.7 | 1.7 | 152 | 28 |
| 3k86 | apo | O87008 | TftC chlorophenol-4-monooxygenase component 1 | tftC | *Burkholderia cepacia* | 23.4 | 1.8 | 158 | 27 |
| 3k88 | FAD/NADH | O87008 | TftC chlorophenol-4-monooxygenase component 1 | tftC | *Burkholderia cepacia* | 23.3 | 1.8 | 158 | 27 |
| 3k87 | FAD | O87008 | TftC chlorophenol-4-monooxygenase component 1 | tftC | *Burkholderia cepacia* | 23.5 | 1.9 | 159 | 26 |
| 2ed4 | FAD/NAD | Q5SJP7 | HpaC 4-hydroxyphenylacetate 3-monooxygenase, reductase component | TTHA0961 | *Thermus thermophilus (strain HB8)* | 21.8 | 1.9 | 145 | 33 |
| 2ecu | apo | Q5SJP7 | HpaC 4-hydroxyphenylacetate 3-monooxygenase, reductase component | TTHA0961 | *Thermus thermophilus (strain HB8)* | 21.7 | 1.9 | 145 | 33 |
| 2ecr | apo | Q5SJP7 | HpaC 4-hydroxyphenylacetate 3-monooxygenase, reductase component | TTHA0961 | *Thermus thermophilus (strain HB8)* | 21.7 | 1.9 | 145 | 33 |
| 4f07 | FAD | O33495 | Styrene monooxygenase component 2 | styB | *Pseudomonas sp. Y2* | 21.5 | 1.9 | 146 | 32 |
| 4xj2 | FMN | A1B5I2 | FerA Flavin reductase domain protein, FMN-binding protein | Pden_2689 | *Paracoccus denitrificans* | 21.2 | 1.9 | 153 | 38 |
| 4xhy | apo | A1B5I2 | Flavin reductase domain protein, FMN-binding protein | Pden_2689 | *Paracoccus denitrificans* | 21.1 | 1.9 | 153 | 38 |
| 3cb0 | FMN | Q8YHT7 | CobR corrin reductase | BMEI0709 | *Brucella melitensis* | 23.2 | 2 | 159 | 29 |
| 4ira | FAD | Q8YHT7 | CobR corrin reductase | BMEI0709 | *Brucella melitensis* | 23.2 | 2 | 159 | 29 |
| 2qck | apo | A0JVA7 | Flavin reductase domain protein, FMN-binding protein | Arth_1583 | *Arthrobacter sp. (strain FB24)* | 21.6 | 2 | 152 | 22 |
| 2d37 | FMN/NAD | Q974C9 | HpaC Putative phenol hydroxylase small component | pheA2 | *Sulfolobus tokodaii* | 21.4 | 2 | 151 | 26 |
| 2d38 | FMN: NADPH | Q974C9 | HpaC Putative phenol hydroxylase small component | pheA2 | *Sulfolobus tokodaii* | 21.2 | 2 | 151 | 26 |
| 2d36 | FMN | Q974C9 | HpaC Putative phenol hydroxylase small component | pheA2 | *Sulfolobus tokodaii* | 20.8 | 2.1 | 152 | 26 |
| 1wgb | apo | Q72LK7 | Probable flavoprotein | TT_C0052 | *Thermus thermophilus* | 20 | 2.5 | 151 | 24 |
| 1yoa | FAD/FMN | Q5SL73 | Probable flavoprotein | TTHA0420 | *Thermus thermophilus HB8* | 19.9 | 2.5 | 151 | 24 |

**Table S3.** Nucleotide and amino acid sequences used in this study.

| | |
|---|---|
| BorF amino acid sequence (Uniprot M9QXS1). | MEGSVNGSQRGNGSQRERVPEPGAGPTTDL<br>LRDSRSLRGIFSSFATGVTVVTVGGDSPHA<br>MTANSFTSVSLDPPLILVCVECDAAMHGSL<br>LEVGSFGVSVLAADQQHVALLYANRWRPRD<br>PTQFDRPGWARGARTGAPLARGALAWFECA<br>LWRAYDAGDHSIFVGRLLTAERHDRRDALV<br>YHSGQFRGLPDRAPVE |
| *borF* codon-optimized nucleotide sequence (GenBank MW847680) | ATGGAAGGTAGCGTTAATGGTAGCCAGCGT<br>GGTAATGGTTCACAGCGTGAACGTGTGCCG<br>GAACCGGGTGCAGGTCCGACCACCGATCTG<br>CTGCGTGATAGCCGTAGTCTGCGTGGTATT<br>TTTAGCAGCTTTGCAACCGGTGTTACCGTT<br>GTGACCGTTGGTGGTGATAGTCCGCATGCA<br>ATGACCGCAAATAGCTTTACCAGCGTTAGC<br>CTGGATCCGCCTCTGATTCTGGTTTGTGTT<br>GAATGTGATGCAGCAATGCATGGTAGCCTG<br>CTGGAAGTTGGTAGCTTTGGTGTTAGCGTT<br>CTGGCAGCCGATCAGCAGCATGTTGCACTG<br>CTGTATGCAAATCGTTGGCGTCCGCGTGAT<br>CCGACCCAGTTTGATCGTCCGGGTTGGGCA<br>CGTGGTGCACGTACAGGTGCACCGCTGGCT<br>CGTGGTGCCCTGGCATGGTTTGAATGTGCA<br>CTGTGGCGTGCCTATGATGCCGGTGATCAT<br>AGCATTTTTGTTGGTCGTCTGCTGACCGCA<br>GAACGTCATGATCGTCGTGATGCACTGGTT<br>TATCATAGCGGTCAGTTTCGTGGTCTGCCG<br>GATCGTGCACCGGTTGAATAACTCGAG |
